# Pediatric Brainstem Encephalitis Outbreak Investigation with Metagenomic Next-Generation Sequencing

**DOI:** 10.1101/414979

**Authors:** Kristoffer E. Leon, Didac Casas-Alba, Akshaya Ramesh, Lillian M. Khan, Cristian Launes, Hannah A. Sample, Kelsey C. Zorn, Ana Valero-Rello, Charles Langelier, Carmen Muñoz-Almagro, Joseph L. DeRisi, Michael R. Wilson

## Abstract

In 2016, Catalonia experienced a pediatric brainstem encephalitis outbreak caused by enterovirus A71 (EV-A71). Conventional testing identified EV in peripheral body sites, but EV was rarely identified in cerebrospinal fluid (CSF). RNA was extracted from CSF (n=20), plasma (n=9), stool (n=15) and nasopharyngeal samples (n=16) from 10 children with brainstem encephalitis or encephalomyelitis and 10 contemporaneous pediatric controls with presumed viral meningitis or encephalitis. Unbiased complementary DNA libraries were sequenced, and microbial pathogens were identified using a custom bioinformatics pipeline. Full-length virus genomes were assembled for phylogenetic analyses. Metagenomic next-generation sequencing (mNGS) was concordant with qRT-PCR for all samples positive by PCR (n=25). In virus-negative samples (n=35), mNGS detected virus in 28.6% (n=10), including 5 CSF samples. mNGS co-detected EV-A71 and another EV in 5 patients. Overall, mNGS increased the proportion of EV-positive samples from 42% (25/60) to 57% (34/60) (McNemar’s test; p-value = 0.0077). For CSF, mNGS doubled the number of pathogen-positive samples (McNemar’s test; p-value = 0.074). Using phylogenetic analysis, the outbreak EV-A71 clustered with a neuroinvasive German EV-A71 isolate. Brainstem encephalitis specific, non-synonymous EV-A71 single nucleotide variants were not identified. mNGS demonstrated 100% concordance with clinical qRT-PCR of EV-related brainstem encephalitis and significantly increased the detection of enteroviruses. Our findings increase the probability that neurologic complications observed were virus-induced rather than para-infectious. A comprehensive genomic analysis confirmed that the EV-A71 outbreak strain was closely related to a neuroinvasive German EV-A71 isolate. There were no clear-cut viral genomic differences that discriminated between patients with differing neurologic phenotypes.

## Introduction

In early 2016, an outbreak of enterovirus A71 (EV-A71) in Catalonia caused more than 100 pediatric cases of neurological disease, ranging from aseptic meningitis to brainstem encephalitis with or without myelitis (1). Virological studies of peripheral body fluids (excluding cerebrospinal fluid [CSF]) identified EV-A71 in almost all patients, but other EVs were also found by EV quantitative reverse transcription polymerase chain reaction (qRT-PCR) during the outbreak, including echovirus-30 (E-30), coxsackievirus (CV)-B1, and CV-A10. The EV-A71 strain was subtyped as subgenogroup C1, and phylogenetic analyses showed it was closely related to an EV-A71 strain associated with a 2015 case of brainstem encephalitis (2, 3). Similar outbreaks occurred in France and Denmark around the same time period (4, 5). EV-A71 was identified from the CSF of 0.02% of patients in the German study, 3% in the Danish study, and 14% in the French study.

Despite this information, questions remained about the Catalonia outbreak. In particular, the EV-A71 subgenotype was only identified by standard testing in the CSF of 11% of patients, all of whom had aseptic meningitis (1, 6), raising the possibility that although almost all the children had a documented systemic EV-A71 infection, their neurological sequelae may have been caused by a para-infectious mechanism or by an unidentified co-infection. In addition, standard qRT-PCR assays only recovered a small segment of the EV-A71 genome, limiting the ability to assess for mutations in the virus genome that may have conferred increased neurovirulence.

Metagenomic next-generation sequencing (mNGS) of CSF is an assay that can simultaneously identify a broad range of infectious agents – viruses, fungi, bacteria and parasites – in patients with neurological symptoms. As opposed to traditional pathogen-specific PCR assays which amplify only limited regions of a microbe’s genome, the entire genome of a pathogen can often be rapidly surveyed with mNGS, making it possible to identify genomic changes in the virus that may correlate with increased neurovirulence or reveal strain divergence (7). Here, we deployed unbiased mNGS of CSF, nasopharyngeal (NP) samples, plasma and stool obtained from affected children during the Catalonia brainstem encephalitis outbreak to screen for additional pathogens (including co-infections) and to compare EV genomes in patients with brainstem encephalitis to patients with more benign neurologic disease (i.e. meningitis with or without encephalitis with self-limited and short-lasting symptoms).

## Methods

### Cohort

The first 10 cases diagnosed with brainstem encephalitis or encephalomyelitis with available residual specimen were included as cases, and the first 10 patients with aseptic meningitis or uncomplicated encephalitis diagnosed during the same period were selected as controls. All of these children were admitted to a tertiary pediatric hospital (Hospital Sant Joan de Deu, University of Barcelona) from April to June 2016. Hospital Sant Joan de Deu is a 300-bed tertiary care hospital for high-complexity patients across a catchment area with a pediatric population of ∼300,000 and has participated in a Spanish EV molecular surveillance network since 2010.

EV-related neurological disease was defined as the detection of EV in any sample in the absence of other causes. The clinical diagnostic approach during the outbreak was previously described (1). The World Health Organization’s Guide to Clinical Management and Public Health Response for Hand-Foot-and-Mouth Disease case definitions were used to assign cases and controls, with case definitions defined in Table 1 (8).

**Table 1.**
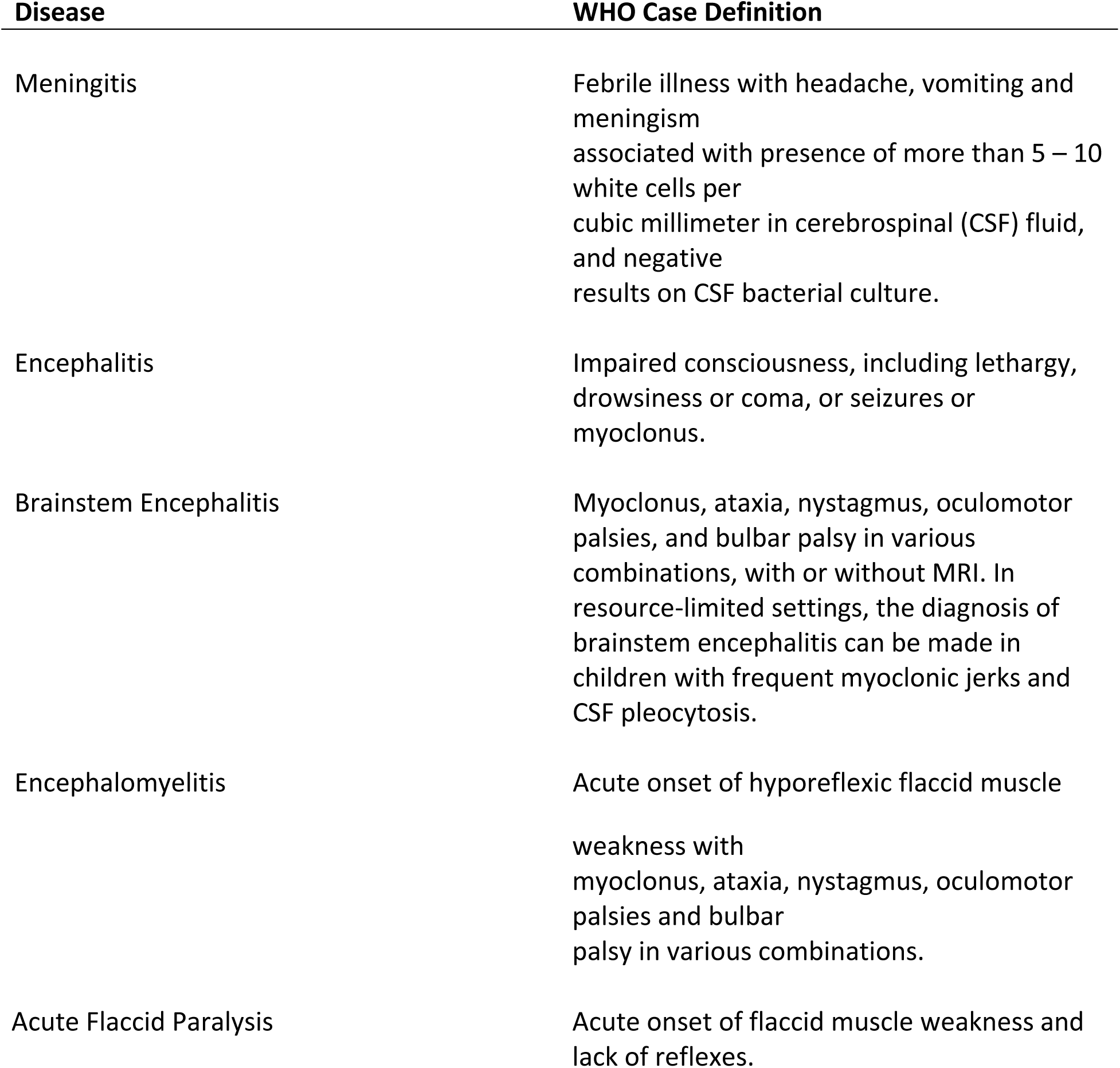
Description of WHO case definitions. *Reproduced from [8].*

Patient demographics and clinical syndromes are described in Table 2. De-identified samples from brainstem encephalitis cases (n=10) were obtained including CSF (n=10), plasma (n=7), NP samples (n=9) and stool (n=9). In addition, de-identified samples from controls (n=10) including CSF (n=10), plasma (n=2), NP samples (n=7), and stool (n=6) were transferred to the University of California, San Francisco for research-based mNGS testing (Supplementary Figure 1). Among these 60 samples, 35 were EV-negative by clinical Pan-EV qRT-PCR. The Pan-EV qRT-PCR is a one-step RT-PCR amplification with EV primers and probes targeted at a conserved region of the 5’ untranslated region, with a limit of detection of 592 genomic equivalents/mL of CSF (9, 10). Patients 1, 3-7 and 9-10, defined as cases, were also negative by the BioFire FilmArray Meningitis/Encephalitis panel (11). All EV-positive patients (by clinical PCR assay) were genotyped at the Enterovirus Unit of the Spanish National Centre for Microbiology using a RT-nested PCR in the 3’-VP1 region specific for species EV-A, B and C, and Sanger sequenced according to a previously described procedure (results reported in Supplemental Figure 1) (12).

**Table 2.** Summary of patient group demographics and clinical data.

|  | Cases<br>Total=10 | Controls<br>Total=10 |
| --- | --- | --- |
| Mean age (months)<sup>a</sup> | 22.7 (18.1-31.2) | 10.8 (0.9-37.5) |
| Sex (male) | 4 | 6 |
| **Systemic symptoms** |  |  |
| Fever | 10 | 10 |
| Vomiting | 6 | 4 |
| Diarrhea | 2 | 1 |
| Exanthema | 5 | 3 |
| Enanthema | 8 | 2 |
| **Neurologic symptoms** |  |  |
| Meningismus | 2 | 3 |
| Irritability | 2 | 2 |
| Lethargy | 8 | 4 |
| Headache | 1 | 2 |
| Myoclonic jerks | 6 | 0 |
| Tremor | 5 | 0 |
| Ataxia | 9 | 0 |
| Paresis | 3 | 0 |
| Nystagmus and/or<br>strabismus | 1 | 0 |
| Bulbar palsy | 2 | 0 |
| Medullary symptoms | 3 | 0 |
| **WHO clinical classification** |  |  |
| Meningitis | 0 | 7 |
| Encephalitis | 0 | 3 |
| Brainstem encephalitis | 8 | 0 |
| Encephalomyelitis | 2 | 0 |
| **EV results by Clinical Pan-EV qRT-PCR (positive/total)** |  |  |
| CSF | 0/10 | 5/10 |
| Plasma | 0/7 | 0/2 |
| Nasopharyngeal sample | 8/9 | 3/7 |
| Stool | 5/9 | 4/6 |

<sup>a</sup> Median (interquartile range).

**Figure 1:**
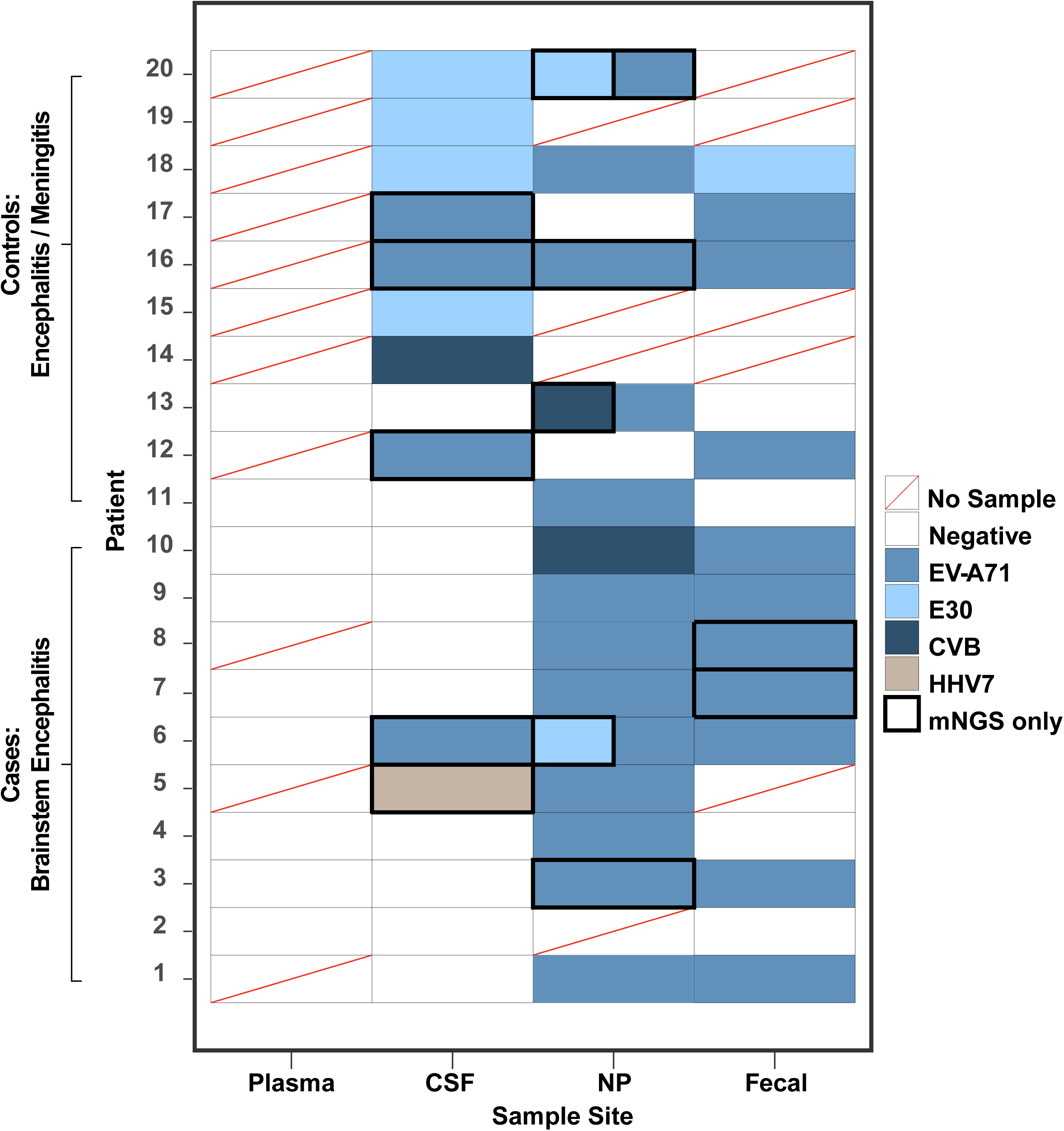
Summary of mNGS diagnostics: concordance and improvement over traditional clinical testing. Unless outlined by a thick, black border, results are concordant. In Patients 3 and 13, EV-A71 was detected by both mNGS and qRT-PCR in the NP samples, however mNGS also detected E30 and CVB respectively. CSF = Cerebrospinal Fluid, NP = Nasopharyngeal, EV-A71 = Enterovirus A71, E30 = Echovirus E30, CVB = Coxsackievirus B, HHV7 = Human Herpes Virus 7

### Metagenomic Sequencing Library Preparation

Samples were received frozen and stored at −80°C. RNA isolation from CSF, serum and NP samples was performed using the RNeasy Micro kit (Qiagen, Germantown, MD). Serum, stool and NP samples were first homogenized with OMNI-International’s 2.8mm ceramic bead kit and the Tissuelyzer II (Qiagen) for 5 min at 15Hz. Stool was extracted using the RNeasy PowerMicrobiome Kit (Qiagen) on a Qiacube (Qiagen). Sequencing libraries were prepared with New England Biolabs’ (NEB; Ipswich, MA) NEBNext RNA First Strand Synthesis Module (E7525) and NEBNext Ultra Directional RNA Second Strand Synthesis Module (E7550) to generate complementary DNA (cDNA). The cDNA was converted to Illumina (San Diego, CA) libraries using the NEBNext Ultra II DNA library preparation kit (E7645) and amplified with 11 PCR cycles. Pre-amplification steps were automated on a Beckman-Coulter Biomek liquid handling robot. The libraries were subjected to Depletion of Abundant Sequences by Hybridization (DASH), described previously, to remove human mitochondrial cDNA (13). The pooled library was size-selected using Ampure beads, and concentration and quality was determined using a Fragment Analyzer (Advanced Analytical Technologies, Inc, Ankeny, IA). Samples were sequenced on an Illumina HiSeq 4000 instrument using 140/140 base pair (bp) paired-end sequencing.

### Bioinformatics

Sequences were analyzed using a rapid pathogen identification computational pipeline developed by the DeRisi Laboratory, described previously in detail (14). Sequences that mapped to the EV genus were collected and *de novo* assembled using the Geneious and/or <u>S</u>t. <u>P</u>etersburg genome <u>A</u>ssembler (SPAdes) algorithms (15). Phylogenetic trees were created in Geneious v10.2.3 using a <u>MU</u>ltiple <u>S</u>equence <u>C</u>omparison by <u>L</u>og-<u>E</u>xpectation (MUSCLE) or <u>M</u>ultiple <u>A</u>lignment using <u>F</u>ast <u>F</u>ourier <u>T</u>ransform (MAFFT) alignment algorithm followed by the Geneious Tree Builder tool, using the Neighbor-Joining build method (16, 17). Bootstrap analysis for each tree was performed with 100 replicates. For single nucleotide variant (SNV) analysis, Bowtie2 v2.3.3 was used to map reads to a reference and then analyzed using VarScan v2.3.4 (18, 19). Statistics on the degree of concordance between research-based mNGS results and standard clinical diagnostic testing results were performed using McNemar’s statistical test.

## Results

### Clinical Testing vs mNGS

We obtained an average of 21.8 million (4.53-61.1 million) 140 bp paired-end reads per sample (Supplemental Figure 1). The non-human sequence reads from each sample have been deposited at the National Center for Biotechnology Information (NCBI) Sequence Read Archive, BioProject (pending). The water controls for CSF and NP samples contained no EV reads. The water control for the stool samples had 0.7 EV rpm when it was pooled and sequenced together with many high EV titer stool samples. To differentiate whether the EV reads present in the stool water control stemmed from physical cross contamination or from bioinformatic contamination due to barcode misassignment (that still occurs at very low levels despite a dual index barcoding strategy) (20), we re-sequenced the same stool water control library separate from high EV titer samples. When sequenced separately, we found no EV reads in this water control. This suggested that the small number of EV reads in the first dataset stemmed from barcode misassignment from the high EV titer samples and not from physical cross contamination. In addition, we re-sequenced the patient samples (NP samples from patients 1, 9, 16 and 18) whose EV read abundance fell below the abundance level of the stool water control on the first sequencing run to similarly determine whether the EV reads were present due to barcode misassignment. Unlike the stool water control, when these patient samples were re-sequenced, they retained EV reads, providing evidence that EV was indeed present in these samples.

Of the 10 cases and 10 controls investigated, EVs were detected in CSF, NP and stool samples via mNGS (Figure 1). Overall, mNGS was concordant with the positive qRT-PCR results for EVs (25 of 25). In the samples found to be EV-negative by clinical Pan-EV qRT-PCR (n=35), pathogens were detected in 28.6% (n=10) of the samples. This included the detection of additional pathogens in 5 CSF samples, including 4 cases of EV-A71 and 1 case of human herpesvirus 7 (HHV-7). Overall, EV-A71 was found in 80% of patients (n=16), E-30 in 25% (n=5), CV-B in 15% (n=3) and HHV-7 in the CSF of 1 patient. Furthermore, 5 patients with EV-A71 infection had another EV detected by mNGS that could not be distinguished by the Pan-EV qRT-PCR assay. Three of these patients had 2 different EVs detected in a single sample (Patients 6, 13 and 20). Overall, mNGS increased the proportion of EV-positive samples from 42% (25/60) to 57% (34/60) (McNemar’s test; p-value = 0.0077). For CSF in particular, mNGS doubled the number of pathogen-positive samples from 5 (25%) to 10 (50%) (McNemar’s test; p-value = 0.074).

### Phylogenetics

Six full-length EV-A71, 4 E30 and 1 CV-B virus genomes (22-2,296x average coverage depth, Genbank MH484066-MH484076) were assembled as described in Methods. All 6 EV-A71 genomes were nearly identical to the German neuroinvasive EV-A71 strain (Genbank KX139462.1, 99.3—99.4% nucleotide similarity and 99.7-99.8% amino acid similarity) (Figure 2b). In addition, we corroborated the previously published phylogenetic analysis that used a Sanger sequence of 360 bases in the 3’ end of the VP1 region, by comparing the entire VP1 gene from our full-length EV-A71 genomes to the VP1 sequences from 11 German neuroinvasive EV-A71 strains, 17 neurovirulent Chinese EV-A71 strains, 6 contemporary, neurovirulent African strains and 26 EV-A71 strains associated with hand foot and mouth disease in Spain (21-24).

**Figure 2:**
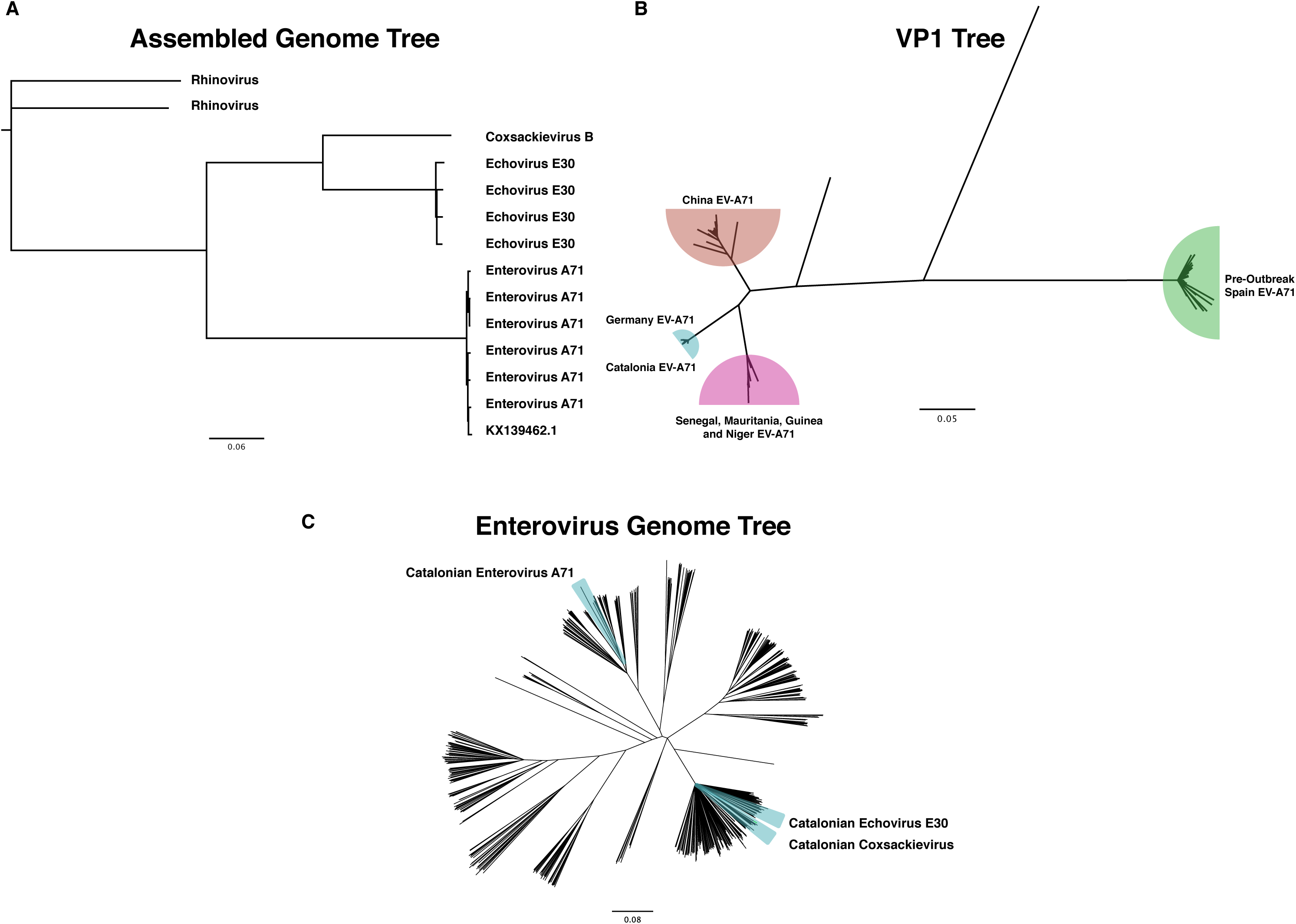
(A) Phylogenetic tree of full length viral genomes for Enterovirus A71, Echovirus E30, Coxsackievirus B and Rhinovirus isolated from CSF, Stool and NP compared to the German neuroinvasive strain. Rhinovirus obtained from patients acts as the root. (B) Confirmation of clinical VP1 testing that the Catalonian EV-A71 Viral Protein 1 (VP1) gene is most closely related to a neuroinfectious German strain. (C) Phylogenetic tree of 545 enterovirus genomes from every species highlighting the relatedness between the enterovirus strains discovered in this outbreak.

Next, 8,841 full length EV genomes were clustered at 95% similarity to create a list of 535 sequences. The EV-A71, E-30 and CV-B strains were aligned to the clustered list using a MAFFT alignment algorithm, followed by phylogenetic analysis. All 3 viruses appropriately clustered within their own species (Figure 2C).

Lastly, SNV analysis was performed on the full-length EV-A71 genomes (3 from patients with brainstem encephalitis, 2 from patients with meningitis and 1 from a patient with encephalitis) to identify SNVs unique to brainstem encephalitis patients. We identified 55 inter-host SNVs that were unique to the EV-A71s associated with brainstem encephalitis. Only 2 of those SNVs were common to all 3 viruses (Supplemental Figure 2). However, both of these SNVs were synonymous mutations located in the 3A and 3D genes, and thus the biological significance of these mutations, if any, is uncertain (25).

## Discussion

The original description of the 2016 pediatric brainstem encephalitis outbreak in Catalonia identified EV-A71 as the likely etiologic agent. However, this conclusion was tempered because 1) there were multiple other co-circulating EVs present during the outbreak, and 2) EV-A71 was not identified in the CSF of the vast majority of patients with brainstem encephalitis. Identifying EVs in peripheral body sites of patients with severe neurologic disease but failing to find it in the CNS mirrors both the recent North American outbreak of acute flaccid myelitis associated with EV-D68 (26) and in an EV-A71 outbreak in the early 2000s (27). As a result, others have hypothesized that the severe neurologic sequelae experienced by children in both outbreaks may have been due to a para-infectious mechanism (or a co-infection) rather than as a direct effect of the virus. Here, we utilized mNGS to further investigate patient samples from the Catalonia outbreak to address these hypotheses.

In contrast to EV-specific qRT-PCR, mNGS identified EV-A71 in the CSF of 4 patients (20%), 1 with brainstem encephalitis, 1 with simple encephalitis, and 2 with meningitis alone. EV-A71 abundance levels in the CSF were very low (0.03-0.55 rpm), and only 3.6% (2/56) of reads mapped to the 5’UTR (the area of the EV genome targeted by the clinical qRT-PCR assay). This may explain the lower sensitivity of the clinical qRT-PCR assay and the BioFire FilmArray panel (1, 6).

With regard to co-infections, we identified HHV-7 in the CSF of 1 patient with brainstem encephalitis who also had EV-A71 identified in the NP sample and stool. There is controversy about the neuro-pathogenicity of HHV-7, although there are a number of reported cases of HHV-7 encephalitis both in immunocompromised and immunocompetent children, including children with brainstem encephalitis (28). In addition, mNGS also identified 5 patients co-infected by 2 different EVs, and 2 of these patients had severe clinical phenotypes, raising the possibility that co-infection may have contributed to disease severity. These findings highlight that in the midst of an outbreak, mNGS can identify patients with alternate or co-infections that produce clinical phenotypes mimicking or contributing to the phenotype associated with the outbreak pathogen.

A limitation of this paper is that, due to sample availability, we did not perform orthogonal confirmation of the mNGS-only virus identifications. As a result, our evaluation of the performance of the mNGS assay is vulnerable to incorporation bias because the gold standard by which we are evaluating its performance includes the mNGS results (29). However, for reasons described above in Results, we are confident that even the low levels of virus detected by mNGS were not due to physical contamination and/or barcode misassignment.

Through mNGS, we were able to recover significantly more genetic information about the EV-A71 outbreak strain, including 6 full-length genomes. This additional information corroborated that the EV-A71 strain was nearly identical to the 2015 neuroinvasive German strain as opposed to the much more common outbreaks of EV-A71-associated brainstem encephalitis seen throughout Asia (Figure 2b-c). Indeed, we found that the strain circulating in Catalonia was more closely related to recently reported contemporary West African strains than Chinese strains (21) (Figure 2b). Despite having full-length genomes, we were unable to identify the presence of any novel or previously known SNVs that might be associated with increased neurovirulence in this limited cohort. Of note, mutations in the 3A (membrane binding protein important in replication) and 3D (the RNA-dependent RNA polymerase) genes have been identified as important regions for adaption to a neuroblastoma cell line (25). In these 2 genes, we identified 1 synonymous SNV in each, common to 3 EV-A71 genomes from brainstem encephalitis patients but not found in meningitis/encephalitis patients. We also searched for a VP1-31G SNV recently reported to be associated with EV-A71 strains with neuroinvasive potential (30). This SNV was not identified in any of the 6 EV-A71 genomes. Despite this, we believe these data will serve as a valuable resource for future studies that seek to identify the genetic determinants of neuroinvasive EV-A71 infections.

While there have been many case reports and small case series documenting the ability of mNGS to detect a variety of pathogens in the CSF (14, 31-33), there have been very few reports on the actual diagnostic yield of mNGS of CSF compared to traditional assays, including pathogen-specific PCR and the BioFire FilmArray. This study demonstrates that in a cohort of patients with a variety of viral CNS infections, a single mNGS assay was concordant with positive pathogen-specific PCR results and indeed, more sensitive for the detection of EV across all sample types, especially in the CSF. Furthermore, identifying EV-A71 in the CSF of 4 additional patients provides circumstantial evidence that this strain was may be neuroinvasive and that a para-infectious etiology is a less likely explanation for the neurological phenotypes observed during the outbreak. Lastly, the rich viral genomic datasets generated by mNGS enabled more sophisticated analyses about the origins of the outbreak strain and the search for possible neurovirulence factors. While the experiments described herein utilized a research-based CSF mNGS assay, a clinically validated CSF mNGS assay with a 3-7 day turnaround time was recently evaluated in a multi-center prospective study and is now clinically available (34). Thus, this type of unbiased diagnostic approach is no longer relegated only to research settings, and its role in individual patient care and public health outbreak investigations will likely expand.

## Acknowledgements

Sandler and William K. Bowes Jr Foundations (MRW, JLD, LMK), Rachleff Family Foundation (MRW), National Institute of Neurological Disorders and Stroke of the National Institutes of Health under award K08NS096117 (MRW) supported this study. This study was partially supported by a grant from the Spanish National Health Institute [grant number PI15CIII-00020] and The European Regional Development Fund (FEDER funds).

We would like to recognize Amy Kistler, Ph.D. for their intellectual discussions, and Dr. Maria Cabrerizo from Enterovirus Unit of the Spanish National Centre for Microbiology for routine EV genotyping of samples. We thank the patients and their families for their participation in the research study.

## Availability of data and material

| Sample | Accession Number |
| --- | --- |
| 4N | MH484066 |
| 5N | MH484067 |
| 6N | MH484068 |
| 11N | MH484069 |
| 16F | MH484070 |
| 17F | MH484071 |
| 15L | MH484072 |
| 18L | MH484073 |
| 19L | MH484074 |
| 20L | MH484075 |

---

**Supplemental Figure 1:** Sequencing results including RPM for each pathogen detected for each patient in each body site.

**Supplemental Figure 2:** Single nucleotide variant map. The red boxes highlight the two common SNVs unique to the brainstem encephalitis patients found in proteins 3A and 3D. The highlighted mutations are 5008T>C and 5938A>G.

